# A method for storing information in DNA with improved dropout tolerance

**DOI:** 10.1101/2023.06.20.545769

**Authors:** Golam Mortuza, Michael D. Tobiason, Kelsey Suyehira, William L. Hughes, Tim Andersen, Reza Zadegan

## Abstract

Storing information in synthetic DNA oligomers is attractive for archival purposes due to the favorable physical density, stability, and energy efficiency of this storage medium. However, issues with this medium sometimes cause dropout (*i*.*e*., loss of oligomers) which may prevent the recovery of stored information. Here, an improved information storage method derived from the existing “DNA Fountain” method is reported. In this work we have developed and experimentally tested a robust algorithm to write digital data in pools of DNA strands by applying a rateless erasure code (*i*.*e*., fountain code), a Reed Solomon code, and an oligomer mapping code. Our new method includes changes to the fountain code, the oligomer mapping code, and the encoding and decoding processes. We have tested and benchmarked our algorithm vs similar algorithms and found that our method increases robustness to dropout, decreases encoding time, and decreases decoding time. The new method was validated *in-vitro* by successfully storing and recovering 105,360 bits of information. The advantages of the new method make it more appropriate for applications where information recovery is critical, where substantial sequence loss is expected, and/or where computational resources are limited. Furthermore, the inclusion of the novel oligomer mapping code enabled us to mitigate errors by restricting sequences of repeated bases and enhance security by eliminating start/stop codons, thus minimizing the risk of interaction with living cells.

## Introduction

Molecules of single stranded deoxyribonucleic acid, referred to as DNA oligomers, are composed of monomer units known as nucleotides (nts). Each nucleotide (nt) typically contains one of four nucleobases (*i*.*e*., adenine, thymine, guanine, or cytosine) and the chemical constitution of a given DNA oligomer can be described by a sequence of these four bases. Several methods for creating synthetic DNA oligomers of arbitrary base-sequence have been reported [1, 2] and it has been recognized that this enables one to store information in the base-sequence of synthetic DNA oligomers [3, 4]. Relative to established information storage technologies, DNA oligomers exhibit highly favorable stability, physical density, and energy efficiency [5–7]. Several methods for storing information in synthetic DNA oligomers have been reported [8–19], including the “DNA Fountain” method demonstrated by Erlich and Zielinski [20]. However, issues with DNA oligomers such as imperfect synthesis, chemical degradation, or inaccurate sequencing may cause dropout (*i*.*e*., a failure to recover DNA oligomers) which sometimes prevents information recovery [21].

## Methods and Materials

Here, the following information storage method was used. At a high level of abstraction (Figure 1A), this new method encodes a digital file into DNA oligomers via a write process and recovers the file from DNA oligomers using a read process. In greater detail (Figure 1B), this method generates binary sequences according to a rateless erasure code (*i*.*e*., a fountain code), extends these binary sequences using a Reed-Solomon error correction code, and then converts these extended binary sequences to base-sequences using an oligomer mapping code. First, a number of oligomers equal to the number of file segments is generated. Next, new oligomers are incrementally generated until the file can be successfully decoded. Once the file has been successfully decoded, additional oligomers are generated until the redundancy (*α*) is greater than or equal to a requested value. Redundancy is calculated according to the following equation:

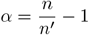

where *n* is the number of oligomers in the current decoding and *n’* is the number of oligomers in the first successful decoding. The processes for generating extended binary sequences and converting them to DNA base-sequences are detailed in the following paragraphs.

**Figure 1:**
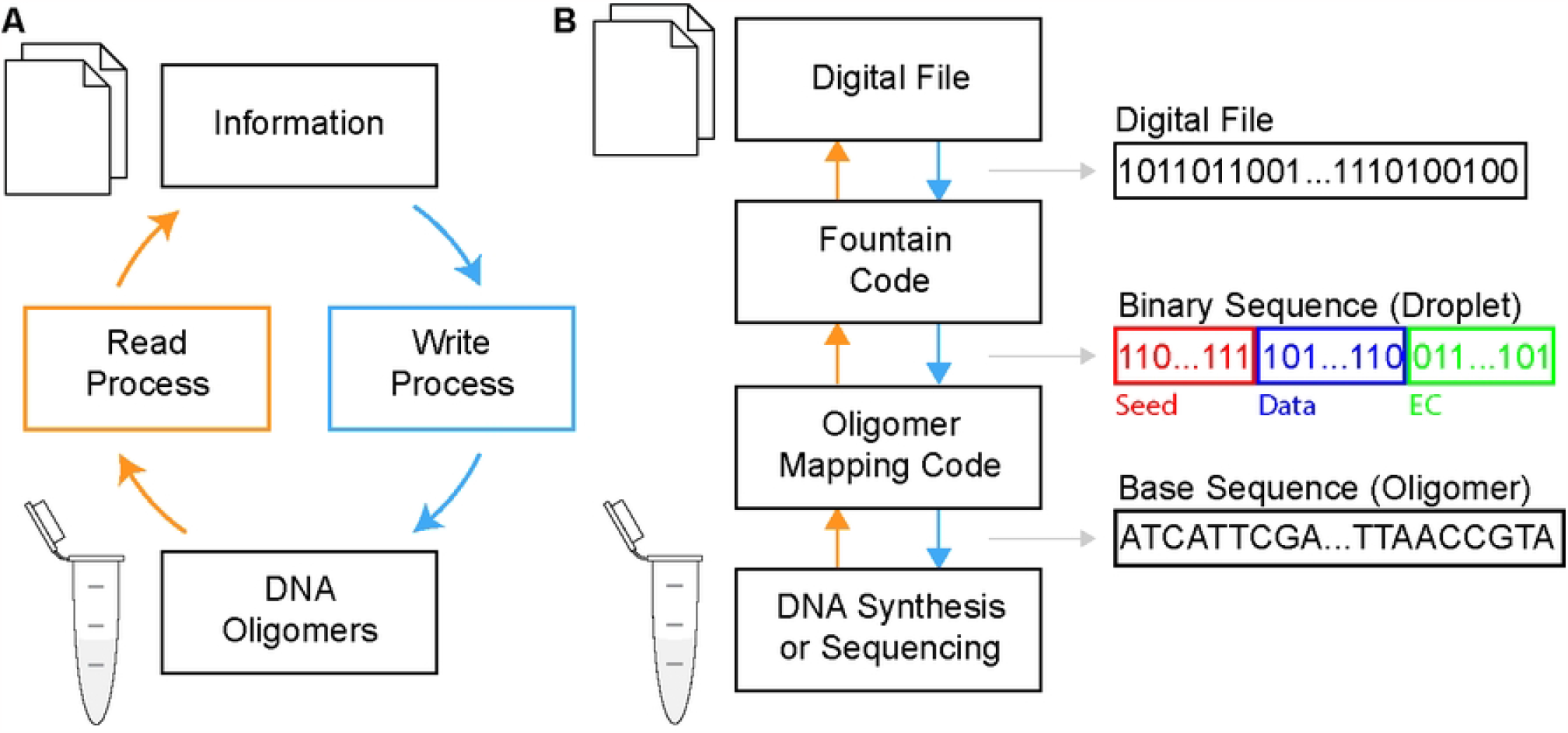
Overview of the new method. (**A**) High level overview. (**B**) Overview of the codes applied in the method. Binary sequences referred to as droplets are generated according to a fountain code. Each droplet includes seed, data, and error correction (EC) bits. Base sequences, referred to as oligomers, which encode a given droplet are generated according to an oligomer mapping code.

The following process was used to generate a binary sequence referred to as a binary droplet. A random integer, referred to as the seed value, was generated using a pseudo-random number generator (PRNG). This seed value was used to initialize a second PRNG, which was used to select segments of the target file to include in the binary droplet. A binary sequence, referred to as the data value, was created by applying an exclusive-or to the file segments specified by the second PRNG. A binary sequence, referred to as the error-correction value, was calculated by applying a Reed-Solomon error correction code [22] to the concatenated seed and data values. The final binary sequence, referred to as the binary droplet, was created by concatenating the seed, data, and error-correction values.

An oligomer (*i*.*e*., sequence of bases) encoding a given binary droplet was generated using the following process. First, the binary sequence is converted to a hexadecimal (hex) sequence. A list of three-base sequences (i.e., codons) corresponding to each hex value is either provided by the user, or the default hex-codon map described below and reported in table 1 is used. A codon is then assigned to the first hex value by selecting the available codon with with highest GC content. This represents the first partial solution of the backtracking algorithm. If GC content is greater than or equal to 50%, the partial solution is extended by assigning the available codon with lowest GC content. If GC content is less than 50%, the partial solution is extended by assigning the available codon with highest GC content. The partial solution is rejected if it contains enough G’s or C’s such that the final sequence could not have acceptable GC content. A partial solution is also rejected it contains sequences of repeated bases larger than a threshold provided by the user. If possible, a rejected partial solution is backtracked by iterating backward through the assigned codons and selecting the next available codon. If no backtrack is possible, a null result is returned. Partial solutions are accepted if a codon was assigned to every hex-value, GC content is acceptable, and no repeat-sequences larger than the threshold are present.

**Table 1.** The hexadecimal-codon map for the new software.

| Hexadecimal | Codons |
| --- | --- |
| 0 | AAC, GAC |
| 1 | AAG, GAG |
| 2 | AGG, GTC, TCT |
| 3 | TCG, CGA |
| 4 | ACT, GCT, TCC |
| 5 | ACC, GCC, CGT |
| 6 | ACG, GCG |
| 7 | AGA, GGA |
| 8 | AGT, GGT |
| 9 | AGC, GGC, CCG |
| a | GAA, CGG |
| b | TAA, CAA |
| c | TAC, CCT, ATC |
| d | TAG, CGC |
| e | TTA, CTA, GTT |
| f | TTC, CTC, GTA |

The default codons corresponding to each hex value are listed in table 1 and were generated using the following considerations [23]. First, of the sixty-four possible three-base codons, four are homopolymers consisting of a single repeated base.

Elimination of these four codons prevent the existence of any homopolymer runs greater than four bases. Of the remaining sixty codons, twelve codons (AAT, ATA, ATT, ATG, CAC, CAT, CAG, CTT, CTG, TAT, TTG, and GTG) are “start codons” used in cellular processes [24] and were removed. Of the remaining forty-eight codons, nine additional codons (ACA, TCA, CCA, GCA, TGA, TGT, TGC, TGG, and GAT) were removed to avoid the creation of “start codons” in the overlap of two adjacent codons. The remaining thirty-nine codons were assigned to hex values based on intuition.

Software implementing the new method was created from the source code provided by Erlich and Zielinski [20]. Relative to this prior method, the following three key updates were made. (1) Oligomers are now generated until the file can be successfully decoded and then additional oligomers are included for redundancy. (2) The binary sequence of each droplet is now converted to a sequence of DNA bases using the new oligomer-mapping code detailed above. (3) Decoding is now done using a breadth-first approach in which a recovered file segment is removed from other binary droplets before any newly recovered segments are processed.

The performance of the new method was compared to that of the prior method using the “first failure test” summarized in Algorithm 1. Briefly described, this test generates a set of oligomers encoding a random file and then randomly removes oligomers until the file can no longer be decoded. This test was performed using the same parameters used in the *in-vitro* validation of the prior “DNA fountain” method [20], which specifically included a target redundancy of 1.07. Four performance metrics were recorded: 1) The dropout fraction at first failure – the fraction of oligomers removed before the file could not be successfully decoded, 2) encoding time – the time required to encode the file, 3) decoding time – the time required to decode the encoded oligomers, and 4) information density – the ratio of bits to the total number of nt in the encoding. Typical performance was estimated using repeated independent trials, and quantified as the median value bounded by the 25th and 75th percentiles. The first failure test was performed for file sizes log-uniformly distributed between 8,192 and 33,554,432 bits (*i*.*e*., 1,024 and 4,194,304 Bytes). The upper limit on file size was dictated by the inconveniently long decoding times exhibited by the prior method.

### Algorithm 1

First Failure Test

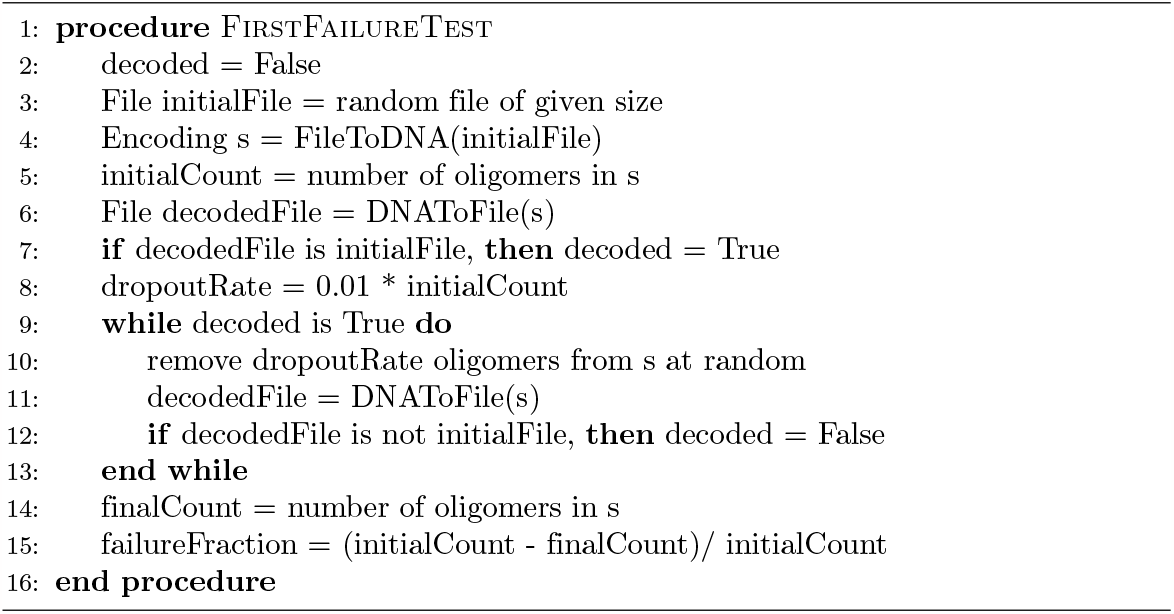

The new method was validated *in-vitro* using the following process. First, the file was encoded into a set of oligomers using the new software. A forward primer (5-ACATCCAACACTCTACGCCC-3) and backward primer (5-GACTACGAGTAGCGCGACGT-3) were attached to each oligomer. The DNA oligomers were then commercially synthesized by Integrated DNA Technologies (1 pmol oPool Oligos). The DNA oligomers were then commercially sequenced by GENEWIZ (Illumina MiSeq sequencing), who was provided no information except the sequence of the primers. The base sequences reported by GENEWIZ were then decoded using the new software.

## Results

Typical *in-silico* performances were estimated using thirty-two independent trials of the “first failure test” and are reported in figure 2. The following notable trends were observed in these results. (1) For all file sizes, the dropout fraction at first failure of the new method was systematically higher than the prior method (figure 2A). At no data point did the middle 50% of the estimates (*i*.*e*. the 25th to 75th percentiles) overlap, indicating this trend was sufficiently resolved using thirty-two trials. (2) For file sizes less than 3 × 10^6^ bits, the new method generated encodings with higher dropout tolerance than requested and the prior code generated encodings which could not be decoded. Both of these trends can be explained by the low probability of generating droplets containing a single file segment, which are critical to the decoding process. (3) The encoding and decoding times of the new method were systematically lower than the prior method (figure 2B and figure 2C). This was attributed to the changes to the decoding process and related code optimizations. (4) The information densities of the new method were systematically lower than the prior method (figure 2D). This trend was attributed primarily to the new oligomer mapping code, and secondarily to the increase number of oligomer generated by the new encoding process.

**Figure 2:**
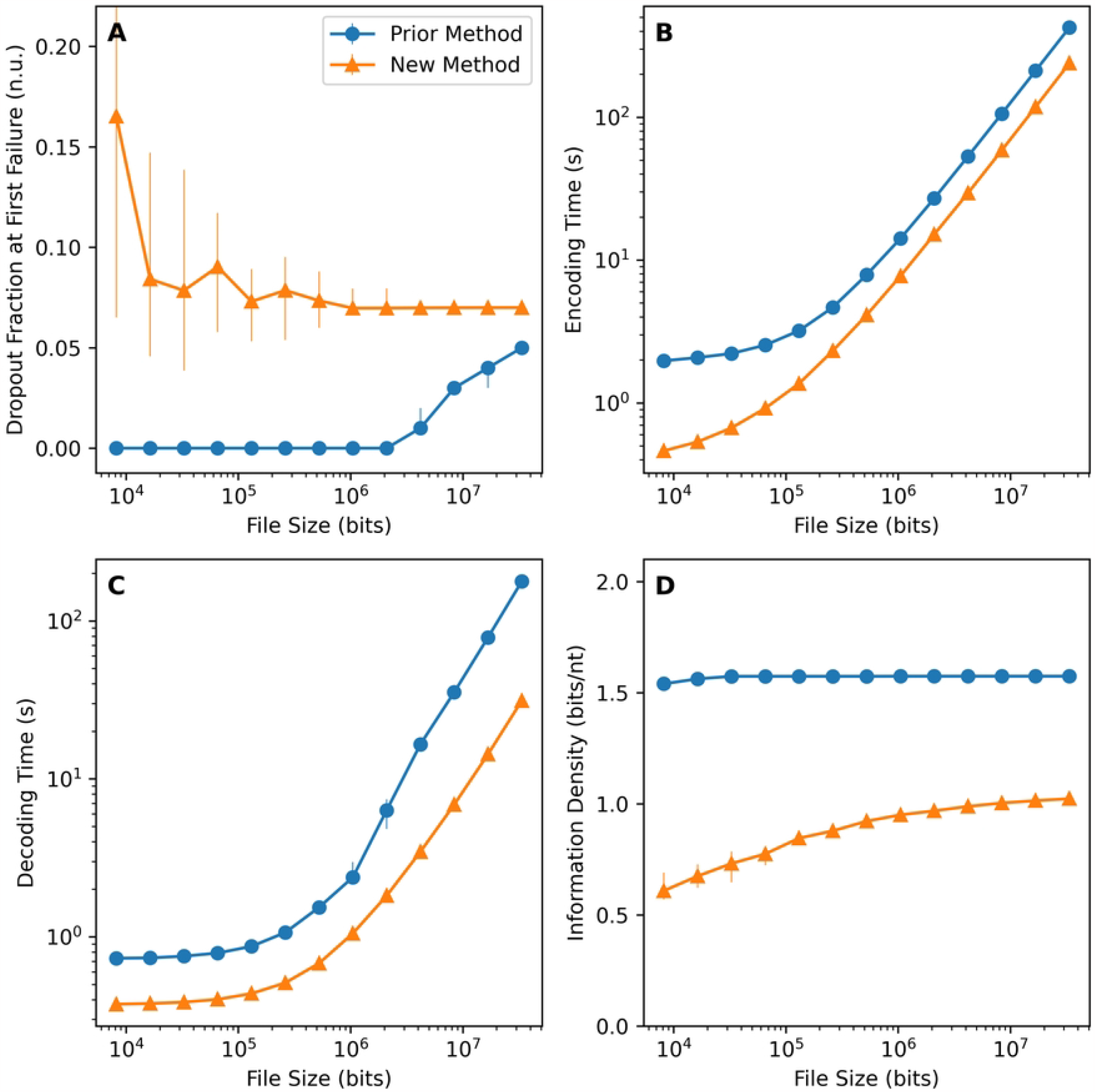
Typical *in-silico* performance of the prior “DNA fountain” method and the new method characterized using the “first failure test”. Each data point represents the median value of thirty-two independent trials and error bars indicate the 25th and 75th percentiles.

The new method was evaluated *in-vitro* by storing and retrieving a 105,360 bit image file (13,170 byte). The image file was successfully retrieved, validating both the new method and new software. The following observations were made regarding this validation. (1) The set of oligomers encoding this image contained 604 oligomers and each oligomer contained 250 nt. This implies a total information density of approximately 0.698 bits/nucleotide. This is 34.9% of the 2 bits/nucleotide limit reported by Church *et al*. [8] and 38.1% of the 1.83 bits/nucleotide Shannon information capacity reported by Erlich and Zielinski [20]. (2) The sequencing process yielded 78 million reads of base-sequences. This included 48 million reads of expected sequences, and 7 million reads of base-sequences which differed from expected sequences by a single base. (3) The 78 million sequencing reads included the base-sequence of 602 of the 604 oligomers from the encoding. This implies a sequence dropout of approximately 0.33%, which is 4 times lower than the approximately 1.3% dropout reported by Erlich and Zielinski [20].

## Discussion

Storing information using the new method was observed to increase dropout tolerance, decrease encoding time, and decrease decoding times. Although our method exhibits a lower information density compared to alternative encoding methods, the advantages of our method make it more appropriate for applications where information recovery is critical, where elevated dropout is expected, and/or where computational resources are limited. The new method additionally contains mechanisms for limiting sequences of repeated bases and eliminating biologically relevant sequences. The former of these is expected to help mitigate synthesis and sequencing errors. The later of these is expected to help mitigate interaction with living cells, preventing an added layer of security against biological interaction.

## Acknowledgments

Not applicable.

## References

1. Kosuri S, Church GM. Large-scale de novo DNA synthesis: technologies and applications. Nature Methods. 2014;11(5):499–507. doi:10.1038/nmeth.2918.

2. Hughes RA, Ellington AD. Synthetic DNA Synthesis and Assembly: Putting the Synthetic in Synthetic Biology. Cold Spring Harbor Perspectives in Biology. 2017;9(1):a023812. doi:10.1101/cshperspect.a023812.

3. Davis J. Microvenus. Art Journal. 1996;doi:10.2307/777811.

4. Feynman RP. There’s plenty of room at the bottom [data storage]. Journal of Microelectromechanical Systems. 1992;1(1):60–66. doi:10.1109/84.128057.

5. Dong Y, Sun F, Ping Z, Ouyang Q, Qian L. DNA storage: research landscape and future prospects. National Science Review. 2020;7(6):1092–1107. doi:10.1093/nsr/nwaa007.

6. Kjær KH, Winther Pedersen M, De Sanctis B, De Cahsan B, Korneliussen TS, Michelsen CS, et al. A 2-million-year-old ecosystem in Greenland uncovered by environmental DNA. Nature. 2022;612(7939):283–291. doi:10.1038/s41586-022-05453-y.

7. Zhirnov V, Zadegan RM, Sandhu GS, Church GM, Hughes WL. Nucleic acid memory. Nature Materials. 2016;15(4):366–370. doi:10.1038/nmat4594.

8. Church GM, Gao Y, Kosuri S. Next-Generation Digital Information Storage in DNA. Science. 2012;337(6102):1628–1628. doi:10.1126/science.1226355.

9. Goldman N, Bertone P, Chen S, Dessimoz C, Leproust EM, Sipos B, et al. Towards practical, high-capacity, low-maintenance information storage in synthesized DNA. Nature. 2013;doi:10.1038/nature11875.

10. Grass RN, Heckel R, Puddu M, Paunescu D, Stark WJ. Robust chemical preservation of digital information on DNA in silica with error-correcting codes. Angewandte Chemie - International Edition. 2015;doi:10.1002/anie.201411378.

11. Tabatabaei Yazdi SMH, Yuan Y, Ma J, Zhao H, Milenkovic O. A Rewritable, Random-Access DNA-Based Storage System. Scientific Reports. 2015;doi:10.1038/srep14138.

12. Blawat M, Gaedke K, Hütter I, Chen XM, Turczyk B, Inverso S, et al. Forward Error Correction for DNA Data Storage. Procedia Computer Science. 2016;80:1011–1022. doi:10.1016/j.procs.2016.05.398.

13. Bornholt J, Lopez R, Carmean DM, Ceze L, Seelig G, Strauss K. Toward a DNA-Based Archival Storage System. IEEE Micro. 2017;doi:10.1109/MM.2017.70.

14. Organick L, Ang SD, Chen YJ, Lopez R, Yekhanin S, Makarychev K, et al. Random access in large-scale DNA data storage. Nature Biotechnology. 2018;doi:10.1038/nbt.4079.

15. Anavy L, Vaknin I, Atar O, Amit R, Yakhini Z. Data storage in DNA with fewer synthesis cycles using composite DNA letters. Nature Biotechnology. 2019;doi:10.1038/s41587-019-0240-x.

16. Lopez R, Chen YJ, Ang SD, Yekhanin S, Makarychev K, Racz MZ, et al. DNA assembly for nanopore data storage readout. Nature Communications. 2019;10(1). doi:10.1038/s41467-019-10978-4.

17. Ping Z, Ma D, Huang X, Chen S, Liu L, Guo F, et al. Carbon-based archiving: current progress and future prospects of DNA-based data storage. GigaScience. 2019;8(6). doi:10.1093/gigascience/giz075.

18. Press WH, Hawkins JA, Jones SK, Schaub JM, Finkelstein IJ. HEDGES error-correcting code for DNA storage corrects indels and allows sequence constraints. Proceedings of the National Academy of Sciences. 2020;117(31):18489–18496. doi:10.1073/pnas.2004821117.

19. Dickinson GD, Mortuza GM, Clay W, Piantanida L, Green CM, Watson C, et al. An alternative approach to nucleic acid memory. Nature Communications. 2021;12(1):2371. doi:10.1038/s41467-021-22277-y.

20. Erlich Y, Zielinski D. DNA Fountain enables a robust and efficient storage architecture. Science. 2017;355(6328):950–954. doi:10.1126/science.aaj2038.

21. Mortuza GM, Guerrero J, Llewellyn S, Tobiason MD, Dickinson GD, Hughes WL, et al. In-vitro validated methods for encoding digital data in deoxyribonucleic acid (DNA). BMC Bioinformatics. 2023;24(1). doi:10.1186/s12859-023-05264-6.

22. Reed IS, Solomon G. Polynomial Codes Over Certain Finite Fields. Journal of the Society for Industrial and Applied Mathematics. 1960;8(2):300–304. doi:10.1137/0108018.

23. Suyehira K. Using DNA For Data Storage: Encoding and Decoding Algorithm Development. Boise State University Theses and Dissertations. 2018;doi:10.18122/td/1500/boisestate.

24. Lobanov AV, Turanov AA, Hatfield DL, Gladyshev VN. Dual functions of codons in the genetic code. Critical Reviews in Biochemistry and Molecular Biology. 2010;45(4):257–265. doi:10.3109/10409231003786094.

